# Neuronal Activity-Dependent Accumulation of Arc in Astrocytes

**DOI:** 10.1101/2020.11.10.376756

**Authors:** Yuheng Jiang, Antonius M.J. VanDongen

## Abstract

The immediate-early gene Arc is a master regulator of synaptic plasticity and plays a critical role in memory consolidation. However, there has not been a comprehensive analysis of the itinerary of Arc protein, linking its function at different subcellular locations with corresponding time points after neuronal network activation. When cultured hippocampal neurons are treated with a combination of pharmacological agents to induce long term potentiation, they express high levels of Arc, allowing to study its spatiotemporal distribution. Our experiments show that neuronal activity-induced Arc expression was not restricted to neurons, but that its spatiotemporal dynamics involved a shift to astrocytes at a later timepoint. Specifically, astrocytic Arc is not due to endogenous transcription, but is dependent on the production of neuronal Arc and accumulates potentially via the recently reported intercellular transfer mechanism through Arc capsids. In conclusion, we found that Arc accumulates within astrocytes in a neuronal activity-dependent manner, which is independent of endogenous astrocytic Arc transcription, therefore highlighting the need to study the purpose of this pool of Arc, especially in learning and memory.

## INTRODUCTION

Activity-dependent gene transcription is a key mechanism that links temporal neuronal activity to down-stream long-lasting effects that eventually cause changes in behaviour. Different classes of activity-dependent immediate-early genes (IEGs) are present in the central nervous system, grouped functionally according to the timeline of expression after neuronal activation (Dragunow 1996), and much work is being done currently to elucidate their mechanisms of action. Arc/Arg3.1 (activity-regulated cytoskeleton-associated protein/activity-regulated gene 3.1) is one activity-dependent IEG that has been shown to be necessary for consolidation of memory in a mouse knock-out model (Plath 2006). Distinct functions of this activity-dependent gene have been demonstrated at various subcellular locations. At the synapse, Arc modulates synaptic plasticity by controlling AMPA receptor endocytosis (Chowdhury 2006), whereas in the nucleus Arc regulates gene transcription of the AMPA receptor gene GluA1 (Korb 2013) and interacts with histone acetyltransferase Tip60 to control acetylation of H4K12, a learning and memory marker that dissipates with aging (Oey 2015). In the endosomal pathway, Arc controls the trafficking of amyloid precursor protein (APP) and presenilin 1 (PSEN1) to the cell membrane, to regulate cleavage of APP (Wu 2011). All these Arc functions are known to be dependent on neuronal activity, although they were studied at different time points and using different mechanisms of neuronal activation. Therefore, the spatiotemporal dynamics of IEG expression profile after neuronal activity is an important functional aspect that should be better studied.

The function of Arc specifically in long-term memory consolidation happens at a late time point, several hours after neuronal activity, corresponding to the late-phase of long-term potentiation (L-LTP). Physiologically relevant expression of phasic Arc has been previously demonstrated to have roles in both fear memory formation (Nakayama 2015) and sleep-dependent consolidation (Seibt 2012). The dynamics of Arc expression seem to vary based on the recent behavioural history of the animal, with high levels of expression for novel contexts and decreased levels for repeated exposures (Guzowski 2006). It is also known that a sustained increase in Arc and short bursts of expression of Arc are elicited with different protocols and may have different effects (Soule 2012, Ramirez-Amaya 2013). Therefore, it seems that although Arc is transiently induced after neuronal activation, the late expression of Arc appears to have a more crucial role in memory consolidation. The timing of expression of Arc and it’s appearance in distinct subcellular compartments are physiologically relevant, for example in novel environment exploration, which results in sustained Arc mRNA and protein expression at 8 hours (Korb 2013, Ramirez-Amaya 2013). Subsequently, it has been shown that lengthening the time of exposure to a novel environment causes Arc protein expression to shift from the cytoplasm to the nucleus (Korb 2013). This indicates that the functional role of Arc in the nucleus is physiologically relevant and specifically timed at the right window for memory consolidation mechanisms to be in play.

IEGs such as Arc are uniquely positioned to transfer information of neuronal activity from neurons to glia. Ongoing cross-talk between glia and neurons has gained increasing attention over the last decade as glia have been shown to not just be a mass of identical cells that simply support neuronal function, but also play a more active role in modulating synaptic plasticity, in a neuronal-activity dependent manner (Araque 1999, Adamsky 2018). Arc has previously been found in astrocytes (Rodriguez 2005, Rodriguez 2008), thus positioning it well for a role to convey information of neuronal activity across cell types. Additionally, Arc has been recently reported to be able to form capsid structures that can mediate intercellular transfer (Pastuzyn 2018), although the functional importance of this form of information transfer remains to be examined. Arc therefore makes an intriguing candidate to study the dynamics of expression after neuronal activation, both within neurons and also possibly in astrocytes.

## MATERIALS AND METHODS

### Neuronal Cell Culture

Hippocampi and cortices from embryonic day 18 Sprague-Dawley rats of either sex were dissected in ice-cold Hanks Balanced Salt Solution (HBSS) and digested using a papain dissociation system (Worthington Biochemical Corporation). The appropriate density of neurons was plated on glass-bottom culture dishes (MatTek, Ashland, MA) that had been coated with poly-D-lysine at 0.1mg/ml (Invitrogen) for 2 hours at 37 degrees. Neurons were cultured in Neurobasal medium supplemented with 2% B-27, 0.5 mM L-glutamine, 10% penicillin/streptomycin, and half of the medium is replaced bi-weekly from day-in-vitro (DIV) 5 onwards. All animal procedures were performed in accordance with the Institutional Animal Care and Use Committee (IACUC) regulations.

### Neuronal Stimulations and Silencing

We used a combination of 4-Aminopyridine (4AP), Bicuculline (Bic), and Forskolin at final concentrations of 100 μM, 50 μM, and 50 μM respectively (4BF) to stimulate synaptic NMDA receptors and network activity and induce Arc expression (Otmakhov 2004, Bloomer 2008, Oey 2015). The drugs were added to the culture medium for two to four hours. Activity silencing was done by adding tetrodotoxin (TTX) at a concentration of 1 μM to the culture medium before treating the culture with the stimulation drugs.

### Conventional Immunofluorescence

Cells were fixed with a solution containing 4% paraformaldehyde (PFA), 4% sucrose, and PBS for 15 minutes at room temperature (RT), blocked with a solution containing 10% goat serum, 2% bovine serum albumin (BSA), and PBS for 1 hour at RT. Primary antibodies were incubated for 1 hour at RT in a dilution buffer containing 1:1 block solution and PBS-Triton X solution at 1:300. Dishes were washed 3 times with PBS-Triton X and incubated with secondary antibodies for 1 hour at RT. Washing was repeated as per the above. The cells were subsequently incubated with 5 μM DAPI for 5 minutes before being imaged. The following primary antibodies are used at the respective dilutions: Mouse-anti-Arc (C7) 1:300 (Santa Cruz Biotechnology Cat# sc-17839, RRID: AB_626696), Rabbit-anti-GFAP 1:1000 (Agilent Cat# N1506, RRID: AB_10013482), Chicken-anti-MAP2 1:1000 (Abcam Cat# ab75713, RRID: AB_1310432), and rabbit-anti-ALIX (Thermo Fisher Scientific Cat# PA5-110566, RRID:AB_2855977). Secondary antibodies conjugated to Alexa-Fluor 488, Alexa-Fluor 568, or Alexa-Fluor 647 (Molecular Probes-Invitrogen) were used in 1:1000 dilution.

### Widefield Imaging

Fluorescence images were obtained using a motorized inverted wide-field epifluorescence microscope (Nikon Eclipse Ti-E), using Nikon 10X Plan Apo objective (N.A. = 0.4) and 40x and 60x Plan-Apo oil objectives, with numerical apertures of 1.35 and 1.49 respectively. Motorized excitation and emission filter wheels (Ludl electronics, Hawthorne, NY) fitted with a DAPI/CFP/YFP/DsRed quad filter set (#86010, Chroma, Rockingham, VT) were used together with filter cubes for DAPI, YFP and TxRed (Chroma) to select specific fluorescence signals. Out-of-focus fluorescence was removed using the AutoQuant deconvolution algorithm (Media Cybernetics).

### Confocal Imaging

Confocal images were acquired in z-series at 60x magnification using an inverted Nikon Ti-E microscope equipped with a Spinning Disc confocal module (CSU-W1, Yokogawa), with a 60X oil Plan-Apo objective (1.49 NA) and a 100X Apo-TIRF objective (1.49 NA). A sCMOS camera (Zyla, Andor) was used to capture the confocal images. Laser lines used were 488 nm (100 mW) for GFP, 515 nm (100 mW) for eYFP and 561 nm (150 mW) for mCherry (Cube lasers, Coherent). Excitation/emission switching was obtained using a dichroic beam splitter (Di01-T405/488/568/647-13 × 15 × 0.5, Semrock) and filter wheels controlled by a MAC6000DC (Ludl). Out-of-focus fluorescence was removed using the AutoQuant deconvolution algorithm (Media Cybernetics).

### Transduction

For calcium imaging, the GCaMP6m construct (pAAV.Syn.GCaMP6m.WPRE.SV40) was a gift from Douglas Kim & GENIE Project (Addgene viral prep #100841-AAV9); http://n2t.net/addgene:100841; RRID:Addgene_100841), and transferred from University of Pennsylvania, Vector Core. It was transduced into the cells at a MOI of 1 × 10^5^ on DIV8. Half of the medium was changed on DIV9 to prevent virus toxicity. Arc shRNA (shArc) and scrambled control Arc shRNA (scrArc) constructs were used as previously described in Leung et. al. (Leung 2019). The Arc-shRNA (shArc) was used in conjunction with adeno-associated virus (AAV9) harbouring the transgenes (concentrations at 1 × 10^13^– 1 × 10^14^ GC/ml range), and synthesized by University of Pennsylvania, Vector Core. On DIV14, cells were transduced at a multiplicity of infection (MOI) of 3 × 10^6^ to ensure a transduction efficiency of around 80%. Half of the medium was changed the next day to prevent virus toxicity.

### Calcium imaging and data analysis

Calcium imaging using widefield microscopy was done on neuronal cultures transduced with GCaMP6. Images were acquired with the Nikon NIS Elements software and subsequently processed with MATLAB, with the BioFormats plugin. Calcium bursts from the whole network were computed based on the total fluorescence change over time. Peaks in fluorescence were detected using the MATLAB function findpeaks. To obtain calcium signals from only the neuronal population, regions of interest (ROIs) were first determined from the DAPI images, and the expression levels of NeuN within the ROIs were quantified (Figure 2.1 B). NeuN positive threshold was determined by mixed population modelling with expression levels from all cells (Figure 2.1 C), and the threshold for negative population was selected to be 3 standard deviations from the population mean. These NeuN positive ROIs are then used to extract fluorescence signals from the calcium movies recorded. All pixels within each ROI are averaged to give a single time course, and calcium transients (ΔF/F) were calculated by subtracting each value with the mean of signals from the lowest 10% of neurons at each time point. This removes background and corrects for photobleaching over time. Spike times were then inferred from ΔF/F using a maximum *a posteriori* (MAP) estimate of the spike train developed by Joshua Vogelstein (Vogelstein, 2009) named fast-oopsi. This implementation was adapted to be used in MATLAB as part of the FluoroSNNAP package developed by Patel and colleagues (Patel et. al., 2015), which is available for download at https://www.seas.upenn.edu/~molneuro/software.html. Firing rates of neurons were calculated as the number of spikes divided by the total recording time.

### Arc expression quantification

Images acquired with the Nikon NIS Elements software are subsequently processed with MATLAB, with the BioFormats plugin. Post processing included a constant background subtraction. To obtain the nuclear expression levels, regions of interest (ROIs) are first determined from the DAPI images, and the Arc expression levels within the ROIs were quantified. Statistical analysis was conducted using GraphPad Prism Software (La Jolla, CA). Data are presented as mean with SEM and statistical significance was set at *P<0.05, **P <0.01, ***P<0.001, ****P<0.0001.

### RNAscope single molecule fluorescent in situ hybridization (smFISH)

We performed smFISH utilizing the RNAscope Fluorescent Multiplex Kit v2 (Cat # 323100 Advanced Cell Diagnostics (ACD), Hayward, California) according to manufacturer’s instructions. Cells were fixed with 4% paraformaldehyde (PFA) for 15 min at room temperature, dehydrated with ethanol and pre-treated with protease IV for 30 min at 40 degree Celsius. Samples were then boiled with Target Retrieval solution for a maximum of 5 mins at 90 degree Celsius. Samples were incubated with commercially available Arc mRNA (Cat # 317071, ACD) and Arc intron (Cat # 432951-C3, ACD) probes. Probes were fluorescently labelled with green (excitation 488 nm), or far red (excitation 647 nm) fluorophores using the Amp 4 Alt A-FL. Subsequently, images were blocked and stained using the immunofluorescence protocol described in the above section to label for GFAP-positive glial cells.

### Conditioned Media Transfer

Neuronal cells are grown in 6-well plates and imaging dishes. Neuronal cultures were treated with 4BF for 4 hours, after which all medium was removed, and replaced with fresh medium for 20 hours. Conditioned medium was harvested and then put onto a fresh neuronal culture (previously untreated). For baseline conditioned medium, no drugs were given, but fresh medium was replaced after 4 hours and incubated for 20 hours before the transfer onto fresh cells cultured in glass-bottom dishes.

### Inhibition of endocytosis using dynasore

Pastuzyn et al. found that endocytosis blockade by application of dynasore reduced the formation of Arc capsids (Pastuzyn 2018). To determine if astrocytic Arc can be reduced via the same mechanism, we pre-treated the neuron for 30 mins with dynasore (Abcam, cat#ab120192) at concentrations ranging from 0.1 to 100 microM, and subsequently subjected them to 4BF for 24 hours.

### Novel environment exposure (NEE) paradigm

C57BL/6 mice were generated from breeding pairs from the colony at the Jackson Laboratory. All animal procedures were performed in accordance with the Institutional Animal Care and Use Committee (IACUC) regulations. Animals from different ages were handled for 3 days before the experiment to familiarise them to the experimenter and to handling in general. The novel environment exposure (NEE) paradigm entails a 42cm by 65cm arena surrounded with a wall of 30 cm, made of translucent plastic. It is enriched with various plastic toys placed at different locations throughout the arena. For the exploration session, mice were picked up from their home cage and placed into the centre of the arena. The mice were anaesthetised and sacrificed via intraperitoneal injection of ketamine (80mg/kg) and xylazine (5mg/kg) mixture and isoflurane inhalation, followed by intracardiac perfusion with phosphate-buffered saline (PB) followed by 4% paraformaldehyde (PFA) in 0.1M PB. Brains were dissected and post-fixed in 4% PFA overnight at 4 degree Celsius and cryopreserved in 30% sucrose in PB until sectioning. Brain sections of 30 μm were made using a sliding microtome (Leica VT1200S, Leica Microsystems, Wetzlar, Germany) chilled with dry ice. The free-floating sections were permeabilized, blocked and stained with antibodies in the same way as cultured neurons as described previously, before being mounted onto slides.

## RESULTS

### Chemical LTP enhances network bursting and increases neuronal firing rate

To achieve long-term potentiation (LTP) in cultured hippocampal neurons, we made use of a combination of 4-aminopyridine, a blocker of presynaptic K_v_1 family K^+^ channels, bicuculline, a γ-aminobutyric acid (GABA) receptor antagonist, together with forskolin, an adenylyl cyclase activator which enhances NMDA-dependent LTP (Otmakhov 2004). This cocktail, subsequently referred to as 4BF, increased synchronised network bursting in hippocampal cultures (**Fig. 1**). The synchronization of the network can be blocked by APV ((2R)-amino-5-phosphonovaleric acid; (2R)-amino-5-phosphonopentanoate), a selective inhibitor of blocker of NMDA receptors, indicating a critical role for NMDA receptors in the synchronous network bursting.

**Figure 1.**
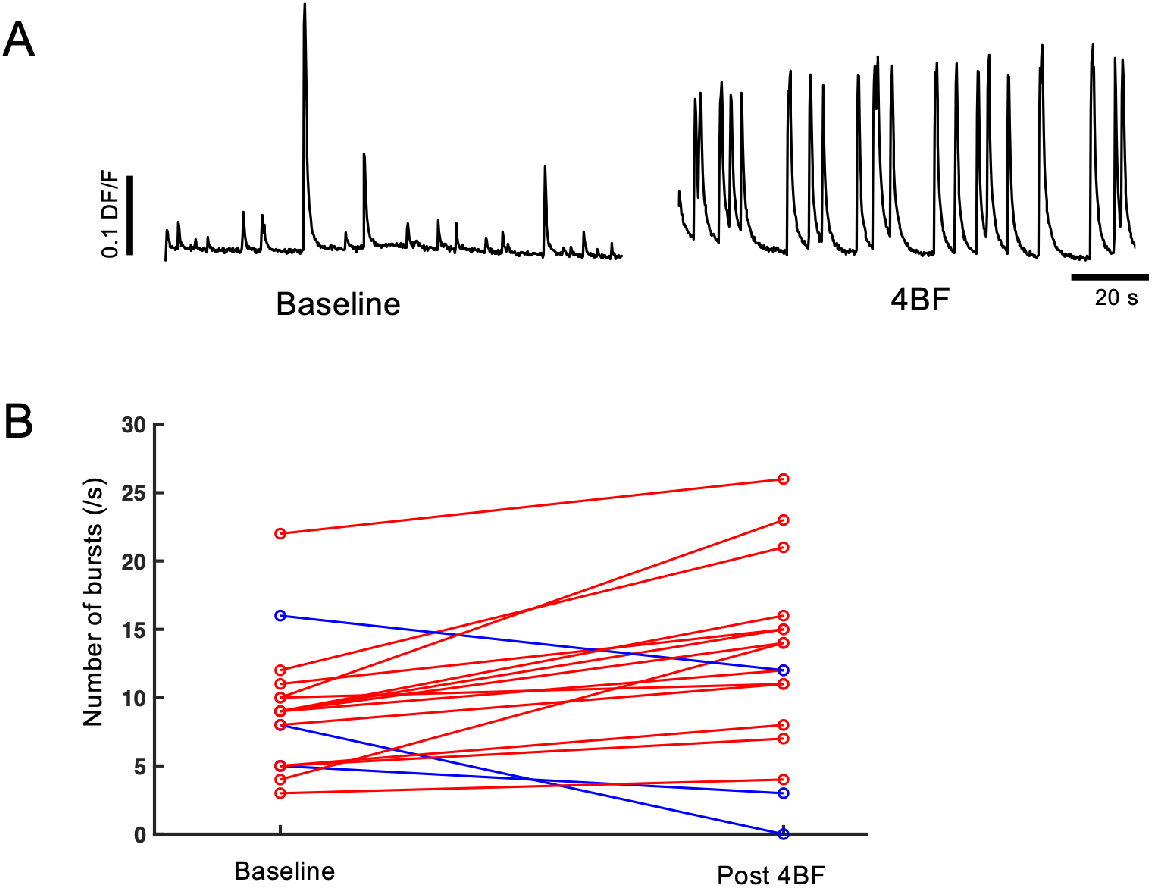
Neuronal cultures exhibit increased network bursting after 4BF treatment. **A**) Representative calcium (dF/F) trace of the whole network before and after 4BF administration, indicating increased network bursts. **B**) Quantification of calcium peaks during 4BF administration and after 4BF removal, as compared to baseline. Increments are marked with red and decrements with blue. Mean difference in calcium bursts = 3.35, SEM = 1.24, n = 17.

### Spatiotemporal dynamics of Arc in neurons and astrocytes

To determine the spatiotemporal distribution of Arc after neuronal activity, the expression of Arc was analysed using immunofluorescence staining at various time points after 4BF induction (**Fig. 2A**). It was found that Arc expression is significantly elevated after 4 hours of drug treatment, with sustained expression up to 24 hours (**Fig. 2B**). To determine the localisation of Arc in different cellular compartments, we analysed its distribution at two time points: 4 hours and 24 hours after 4BF treatment, by specifically quantifying the nuclear as well as cytoplasmic Arc levels. We found that Arc was localized predominantly in the nucleus at 4 hours and shifted to the cytoplasm at 24 hours (**Fig. 2C**).

**Figure 2.**
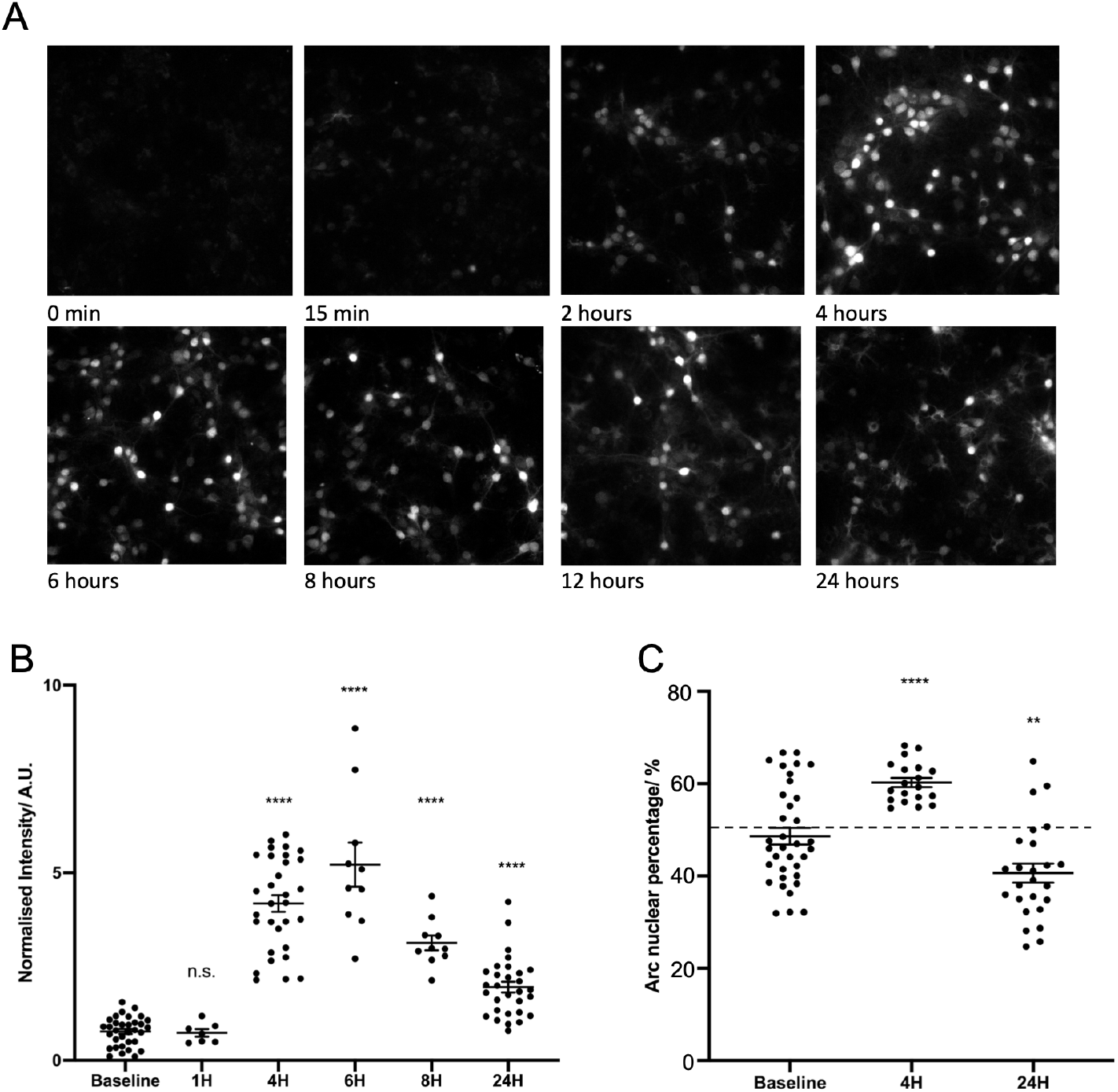
Arc protein expression levels after 4BF. **A**) Timeline of Arc protein expression after pharmacological stimulation by 4AP, bicuculline, and forskolin (4BF). **B**) Arc protein levels are significantly increased from 4 hours to 24 hours after 4BF (one-way ANOVA, F (5, 115) = 65.53, p < 0.0001, post-hoc with Dunnett’s multiple comparison test). A.U. = arbitrary fluorescence units. Quantifications of individual frames from at least 3 biological replicates at each time point, normalised to the baseline fluorescence level from each biological replicate. Baseline (0.77 ± 0.067, n = 33), 1 hour (0.73 ± 0.10, n = 7, p = 0.99), 4 hours (4.18 ± 0.22, n = 31, p < 0.0001), 6 hours (5.22 ± 0.59, n = 10, p < 0.0001), 8 hours (3.13 ± 0.20, n = 10, p < 0.0001) and 24 hours (1.95 ± 0.14, n = 30, p < 0.0001). **C**) At 4 hours, there is significantly more nuclear Arc than at baseline and at 24 hours, there is significantly less nuclear Arc, than at baseline. (one-way ANOVA, F (2, 76) = 23.34, p < 0.0001, post-hoc with Dunnett’s multiple comparison test). A.U. = arbitrary fluorescence units. Quantifications of individual frames from at least 3 biological replicates at each time point. Baseline (48.6% ± 1.8%, n = 35), 4 hours (60.2% ± 1.0%, n = 19, p < 0.0001) and 24 hours (40.6% ± 2.1%, n = 25, p = 0.0036).

To further investigate the distribution of Arc in different cell types, we labelled neurons and astrocytes using the cell type specific markers MAP2 and GFAP, respectively. We found that Arc was present in neurons at 4 hours, and was subsequently found in the GFPA-positive astrocytes (**Fig. 3A**). Taken together, these results suggest that nuclear Arc at 4 hours is found exclusively in neurons, while cytoplasmic Arc at 24 hours is found predominantly within astrocytes. Interestingly, Arc is excluded from the nucleus in astrocytes (**Fig. 3B**). We found that at the late timepoint (24 hours 4BF) Arc was found in close association with GFAP (**Fig. 3C**). This is consistent with previous reports of Arc found in GFAP-positive astrocytes using electron microscopy immunogold labelling technique (Rodriguez 2008), which also reported Arc’s close association with GFAP.

**Figure 3.**
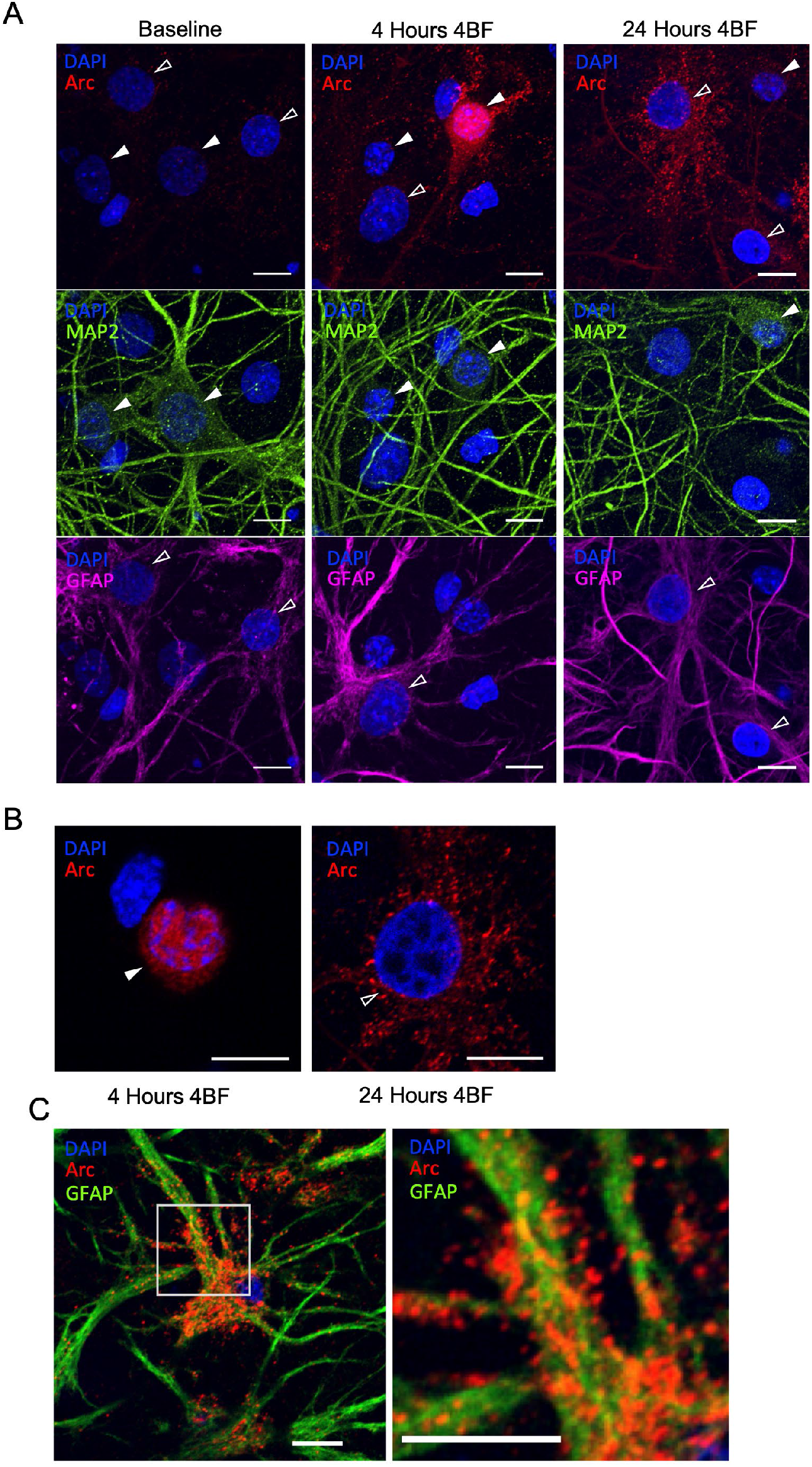
Arc distribution in different cell types. A) Confocal microscopy imaging of Arc expression at the same time points. Co-staining with MAP2 (green), a marker for neurons, and GFAP (magenta), a marker for astrocytes, reveal that Arc expression in the nucleus at 4 hours are in neurons (solid arrowheads) and cytoplasmic expression of Arc at 24 hours is found in astrocytes (open arrowheads). B) Single-slices from z-stack images from 2A at 4 hours and 24 hours, indicating that Arc is nuclear in neurons and excluded from the nucleus in astrocytes. C) Astrocyte with Arc at 24 hours after 4BF stimulation. Insert shows magnified regional views indicating close association between GFAP (green) and Arc (red).

### Neuronal silencing abolishes both neuronal and astrocytic Arc

Tetrodotoxin (TTX) is a potent blocker of voltage-gated Na channels which strongly inhibits neuronal activity. TTX reduces neuronal activity even in the presence of 4BF (**Fig. 4A**). We examined the effect of TTX on Arc expression at different time points and found that Arc expression at both 4 and 24 hours was reduced to baseline levels with neuronal silencing by TTX treatment (**Fig. 4B**). This suggests that prior activity-dependent neuronal expression of Arc may be required for the expression of astrocytic Arc at a later time point.

**Figure 4.**
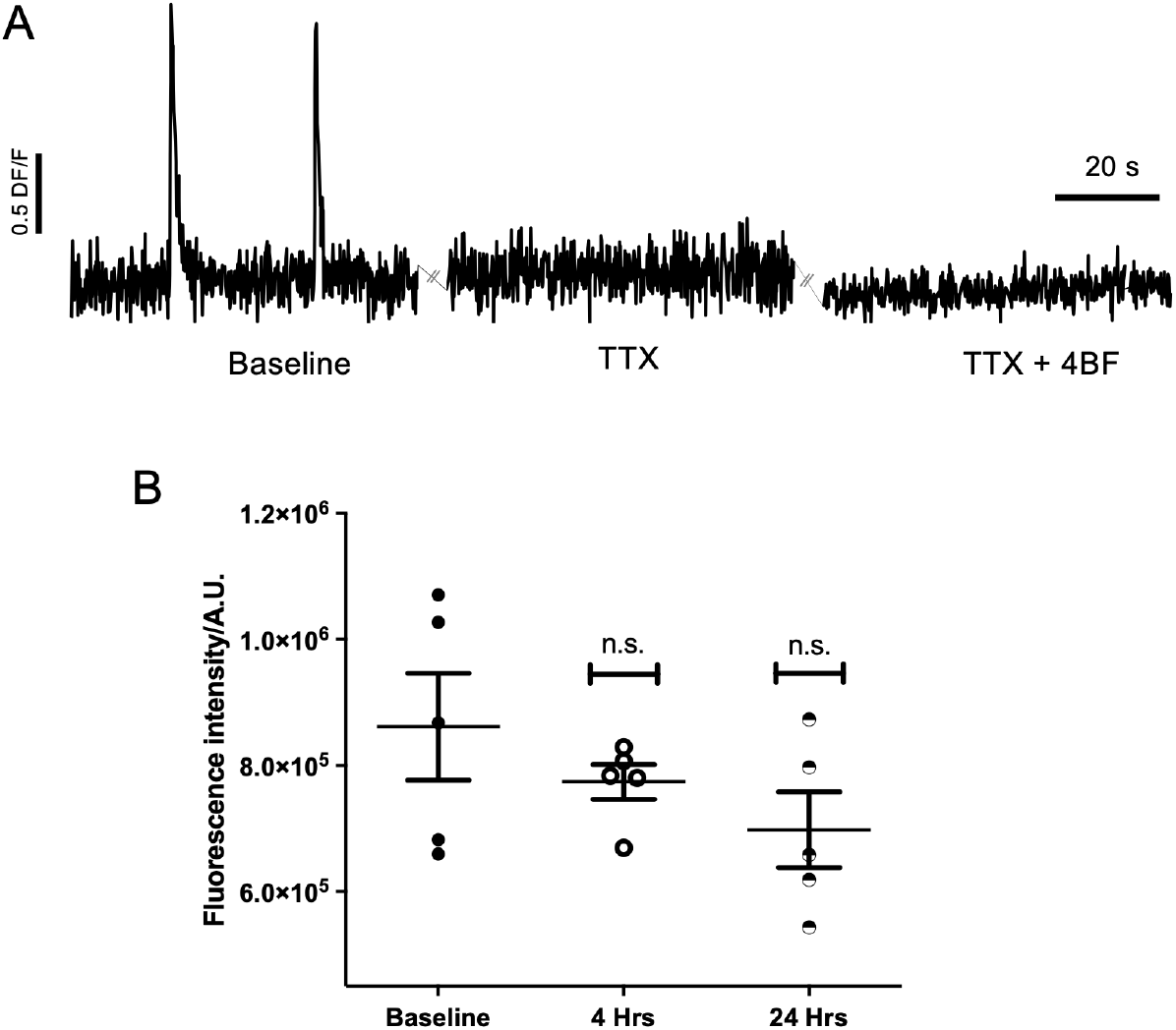
Arc levels after neuronal silencing by TTX. A) TTX silences neuronal cultures in the presence of 4BF, as indicated by the representative calcium trace (ΔF/F). B) Quantification of Arc levels show that 4 hours and 24 hours conditions are comparable with baseline (one-way ANOVA, F (2, 12) = 1.731, p =0.2186, post-hoc with Dunnett’s multiple comparison test). A.U. = arbitrary fluorescence units. Quantifications of individual frames from at least 2 biological replicates at each time point. Baseline (8.6157 ± 0.8488 × 105, n = 5), 4 hours (7.74 ± 0.28 × 105, n = 5, p = 0.6032) and 24 hours (6.9825 ± 0.6010 × 105, n = 5, p = 0.2653).

### Arc is not transcribed in astrocytes

To further investigate the source of Arc within astrocytes, we utilised a single molecule fluorescent in-situ hybridisation (smFISH) probe to specifically target the intronic region of Arc pre-mRNA (Arc-intron RNA). This labels cells that are actively transcribing Arc. Another smFISH probe targeting Arc mRNA was used to identify the cells that contain the mature form of Arc RNA. Arc-intron RNA and Arc mRNA were co-stained with GFAP to determine if there are any foci of transcription or presence of Arc mRNA within astrocytes. We found that Arc-intron RNA and Arc mRNA were found exclusively within non-GFAP-labelled cells at various time points after 4BF (**Fig. 5**). WE found that Arc is actively transcribed 1 hour and 4 hours post 4BF stimulation, and that transcription is drastically reduced at 24 hours after 4BF (**Fig. 5C**). Both the foci of transcription and the pool of Arc mRNA are found exclusively within GFAP-negative cells (at 1-hour post 4BF, with 0 out of 35 cells being GFAP-positive; at 4 hours post 4BF, with 0 out of 12; and at 24 hours post 4BF 0 out of 4). These spatiotemporal results showed that Arc is not transcribed within astrocytes and suggest that astrocytic Arc originates from the neuronal pool.

**Figure 5.**
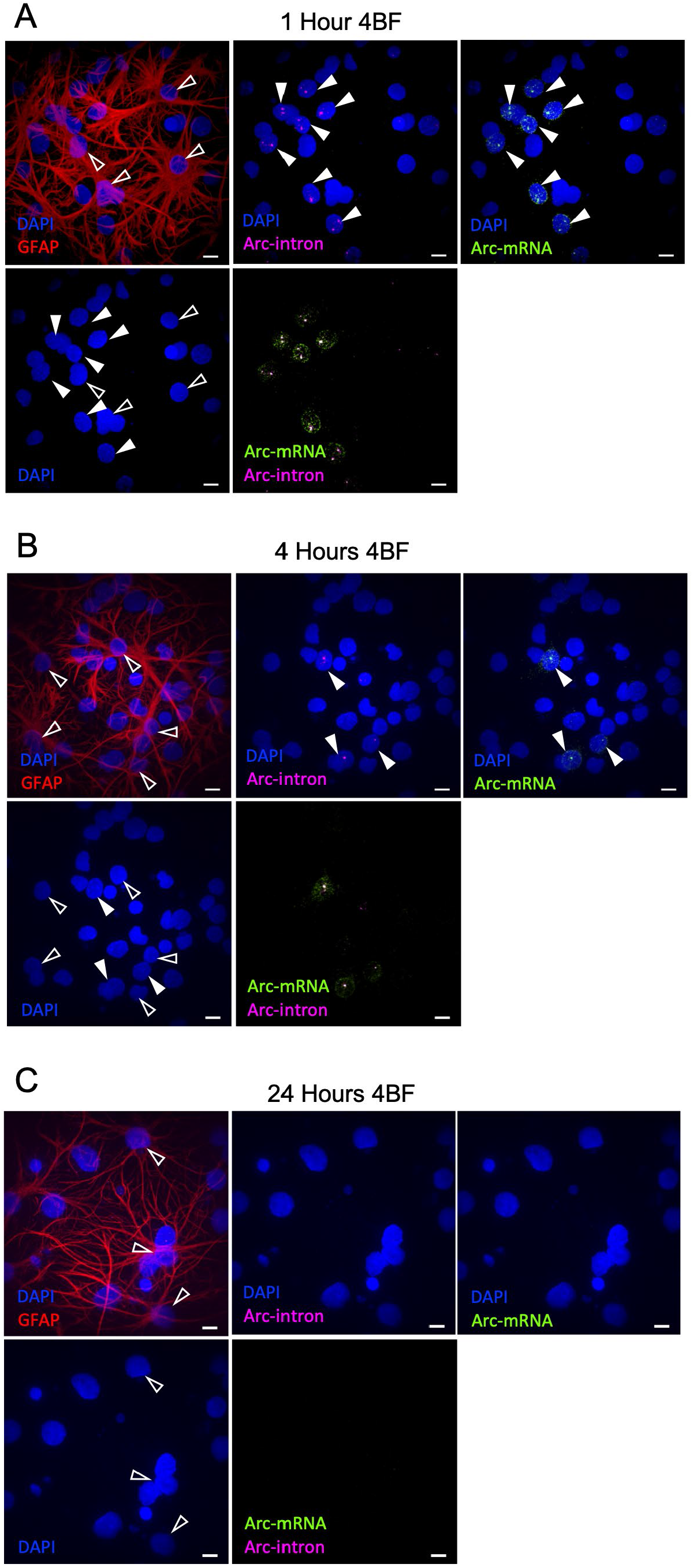
Arc intron and mRNA smFISH. smFISH of Arc-intron probe (magenta) and Arc-mRNA probe (green), labelled together with GFAP (red) and DAPI (blue). (**A**-**C**) 1, 4 and 24 hours after 4BF, showing that Arc-intron foci and Arc-mRNA (solid arrowheads) do not overlap with cells labelled with GFAP (open arrowheads). The lack of Arc-intron and Arc-mRNA signals at 24 hours indicate that Arc transcription is low at this time.

### Astrocytic Arc is dependent on Arc transcription

A previously validated short hairpin RNA (shRNA) construct targeting the coding region of Arc has been shown to be able to suppress Arc induction following 4BF treatment (Leung 2019). Here, we used this shRNA to determine if astrocytic Arc is dependent on Arc transcription. A scrambled version of the Arc shRNA (scrArc) was used as a negative control, to demonstrate that Arc can be successfully induced at 4- and 24-hours post 4BF (**Fig. 6A**). In contrast to the effect of the scrambled control, the expression of Arc was reduced to baseline in the Arc-shRNA knockdown cells at both 4 and 24 hours (**Fig. 6B**). This indicates that Arc mRNA is required for the production of Arc protein, which is then observed first in neurons (in the nucleus) and later in the cytoplasm of astrocytes. Quantification of Arc levels within individual AAV9 transduced cells shows that Arc is induced in the scrambled control, while Arc expression is abolished in Arc-shRNA knockdown conditions (**Fig. 6C**). Taken together with the previously established finding that Arc is not transcribed within astrocytes, this indicates that astrocytic Arc levels may depend on transcription of Arc within neurons.

**Figure 6.**
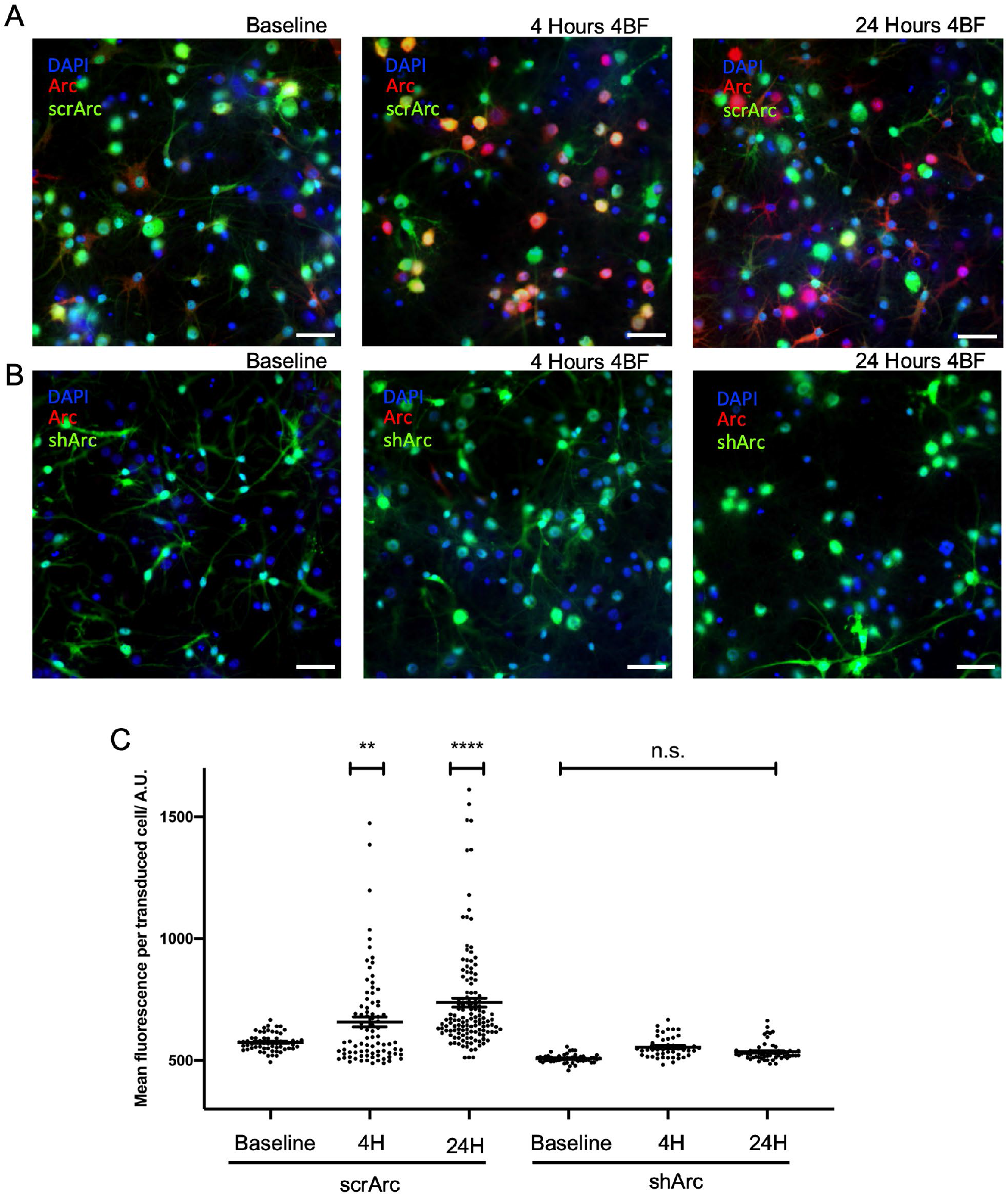
Arc-shRNA knockdown abolished astrocytic Arc. **A**) Knockdown of Arc using shRNA (shArc, bottom row) delivered by an AAV vector abolishes Arc expression in neurons at 4 hours after 4BF and also astrocytic Arc expression at 24 hours. Scrambled Arc shRNA (scrArc, top row) serves as control, where both types of Arc expression remain. **B**) Quantification of results from cultures of both Arc-shRNA and scrambled RNA. Arc levels are quantified in transduced cells, indicating that cells with scramble-RNA knockdown can still express Arc but in cells with Arc-shRNA the levels of Arc within the cells are reduced to baseline. (one-way ANOVA, F (5, 419) = 30.15, p < 0.0001, post-hoc with Dunnett’s multiple comparison test). A.U. = arbitrary fluorescence units. Quantifications of individual frames from at least 3 biological replicates at each time point. Baseline scrArc (574.4 ± 4.6, n = 60), 4 hour scrArc (657.8 ± 20.8, n = 84, p = 0.0039), 24 hours scrArc (737.3 ± 18.2, n = 132, p < 0.0001), Baseline shArc (507.2 ± 2.5, n = 49, p = 0.0717), 4 hour shArc (554.4 ± 6.6, n = 44, p = 0.9345) and 24 hours shArc (535.2 ± 5.0, n = 56, p = 0.4561).

### Astrocytic Arc is dependent upon neuronal Arc production

To further ascertain if astrocytic Arc can accumulate as a result of transfer from neighbouring neurons actively producing Arc, we used conditioned media from prior 4BF-treated cultures on naive cells following the protocol as outlined in **Fig. 7A**. We found that naive cultures receiving conditioned media from active cultures exhibited a substantial increase in both neuronal and astrocytic Arc levels (**Fig. 7B, C**). We further made use of smFISH to determine if Arc present within astrocytes is produced endogenously. WE found that Arc-intron RNA foci were not localised within GFAP-positive cells (**Fig. 7D**), indicating that the increase in Arc levels within astrocytes is not due to its endogenous production. Taken together, these results indicate that astrocytic Arc can be transferred from the Arc pool produced in neuronal cells.

**Figure 7.**
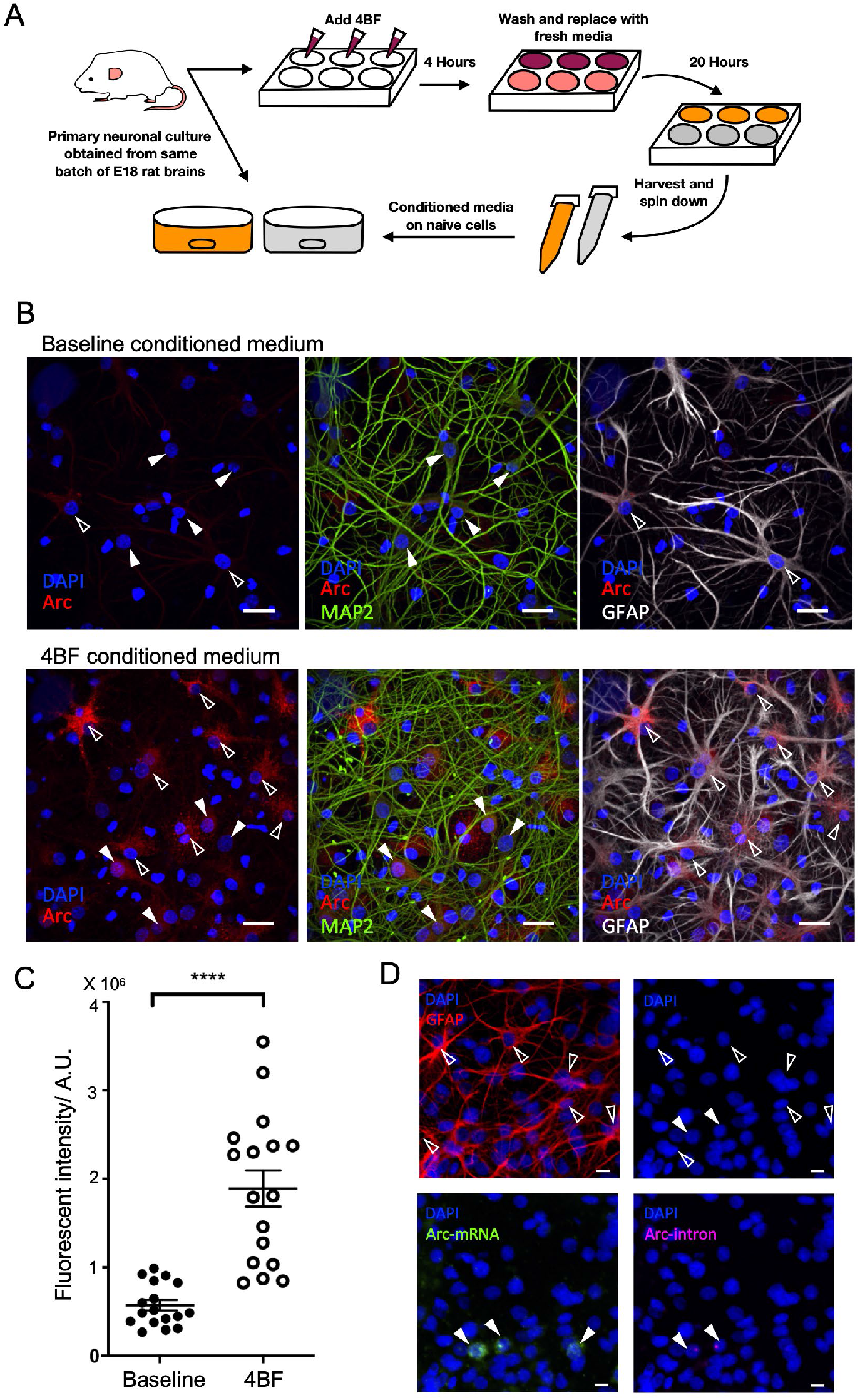
Culture media of active networks can induce astrocytic Arc accumulation. **A)** Schematic diagram indicating the conditioning and transfer of culture medium. Neuronal cultures were treated with 4BF for 4 hours, after which all medium was removed, and replaced with fresh medium for 20 hours. Conditioned medium was harvested and then put onto a fresh neuronal culture (previously untreated). For baseline conditioned medium, no drugs were given, and fresh media were also replaced after 4 hours and incubated for 20 hours before the transfer onto naive cells. **B)**(Bottom row) Arc is induced both in astrocytes (open arrowheads) and neurons (solid arrowheads) after 24 hours incubation from conditioned media not containing 4BF. (Top row) In contrast, conditioned media from cultures with no prior induction do not cause strong arc expression in neurons and astrocytes. **C)** Quantification of Arc transfer indicating that Arc levels are significantly higher with conditioned media from active cultures (activity) as compared with control (baseline conditioned media) (Unpaired t-test, t(32) = 6.189, p < 0.0001) A.U. = arbitrary fluorescence units. Quantifications of individual frames from at least 3 biological replicates at each time point. Baseline conditioned (5.7251 ± 0.5834 × 105, n = 17), 4BF conditioned (18.9138 ± 2.0498 × 105, n = 17). **D)** smFISH with Arc-intron probe (magenta), Arc-mRNA probe (green), GFAP (red) and DAPI (blue), which shows Arc-intron foci and Arc-mRNA within cells not labelled with GFAP (solid arrowheads). Astrocytes labelled with GFAP are labelled with open arrowheads.

### Astrocytic Arc levels are not affected by inhibition of endocytosis by dynasore

Pastuzyn et al. (Pastuzyn 2018) previously showed that capsid Arc colocalizes with ALG-2 interacting protein-X (ALIX), and capsid transfer can be blocked by inhibition of endocytosis using dynasore, a dynamin inhibitor. To determine if astrocytic Arc accumulation is dependent upon endocytosis in the same way, we pre-treated the neurons with dynasore at varying concentrations before subjecting the cultures to 4BF. WE found no significant decrease in Arc levels after dynasore pre-treatment (**Fig. 8A**), indicating that the astrocytic Arc expression may not require endocytosis. WE also showed that astrocytic Arc does not colocalize with ALIX, indicating that they are not associated with endosomes within astrocytes (**Fig. 8B**).

**Figure 8.**
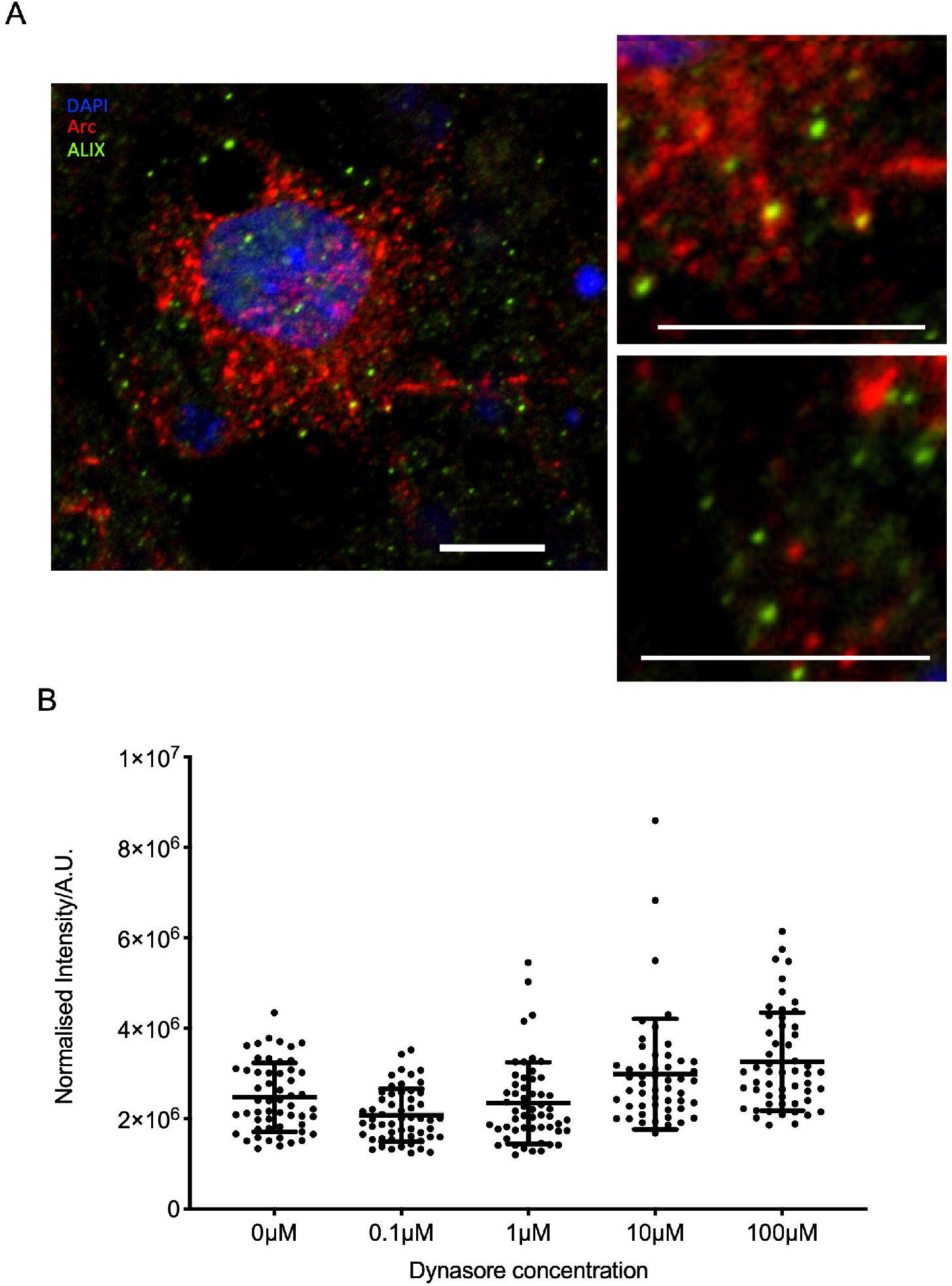
Astrocytic Arc levels in response to dynasore. **A)** Astrocytic Arc levels were not reduced by pre-treatment with Dynasore applied before 4BF at various concentration. **B)** Astrocytic Arc did not co-localise with ALIX. Inserts on the right show magnified views of Arc (red) which do not overlap entirely with ALIX (green).

### Arc induction in vivo using NEE did not reveal astrocytic Arc

To determine if astrocytic Arc can be found in vivo, we used and adapted previously established behavioural protocols of novel environment exploration (NEE, Ramirez-Amaya 2005; (Korb 2013, Ramirez-Amaya 2013). Although Arc was induced in the hippocampal region, the level of induction was low, and cells were sparsely labelled. We did not find any evidence of Arc within astrocytes by examining co-staining with GFAP.

## DISCUSSION

Arc is a well-studied and important player in the processes underlying synaptic plasticity and long-term memory consolidation (Shepherd 2011). However, because methods of inducing neuronal activity vary in the published studies, there has not been any comprehensive analysis to characterize the itinerary of Arc protein, which has been reported to localize to the neuronal soma, dendrites, spines and the nucleus. Its function at the different subcellular locations has also not been linked with the corresponding timing after neuronal activation. We used an *in vitro* system of cultured hippocampal neurons to determine the spatio-temporal distribution of Arc after neuronal network activation, as the cultured neurons are capable of forming networks that produce spontaneous activity and can be treated with a combination of pharmacological agents to induce LTP, resulting subsequently in high expression levels of Arc. We showed that the spatiotemporal dynamics of Arc was not restricted to neurons, but involved a shift to the astrocytes that are part of the network. Specifically, astrocytic Arc is not due to endogenous transcription within the astrocytes, but is dependent on the production of neuronal Arc.

### Nuclear Arc and long-term memory consolidation

We used 4BF to induce LTP in the neuronal network cultures and showed that there is an increase in synchronised bursting as a result of the treatment. As previous studies suggest, Arc is induced in an NMDA receptor-dependent manner (Bloomer 2008). In line with this, we noted low baseline levels of Arc prior to treatment and increased Arc production with increased synchronised network firing. This elevated Arc production is evident at the network level after 4 hours of treatment, at which time Arc mainly localised to the nucleus of neurons. This implies that our protocol may be similar to the behavioural paradigms which causes Arc expression preferentially in the nucleus, for instance that of the novel environment exploration (NEE), as described previously (Korb 2013, Ramirez-Amaya 2013). The four-hour time point at which this expression peaks also corresponds to a later phase in the commonly used LTP protocol, where a high-frequency electrical stimulus is applied to create the long-lasting potentiation. Therefore, this supports the notion that Arc acts at the late phase of LTP via its role in the nucleus, which subsequently contributes to the consolidation of long-term memory. Nuclear Arc is known to associates with beta-spectrin, PML-bodies (Bloomer 2007, Korb 2013), and chromatin remodelling complexes including Tip60 (Wee 2014, Oey 2015) and Amida (Irie 2000). These all have diverse roles in regulation of gene transcription, via both genetic and epigenetic mechanisms, which are processes crucially involved in the consolidation of long-term memory.

According to our results, the Arc levels are sustained in the presence of the stimulus. Although this may not represent a physiological condition, there has been evidence that suggests that sustained transcription occurs long after the stimulus is given in an *in vivo*, behaviourally relevant setting, generating Arc that is preferentially found in the nucleus (Ramirez-Amaya 2013). In a separate behavioural paradigm with fear conditioning, it was found that Arc was expressed in phases, with an initial transient expression immediately after the stimulus followed by a secondary increase in Arc levels after 12 hours (Nakayama 2015). This late phase of Arc is dependent on brain-derived neurotrophic factor (BDNF) and is essential for memory consolidation. Reports also suggest a higher nuclear to cytoplasmic ratio during sleep (Honjoh 2017), and that sleep-dependent consolidation of cortical plasticity is caused by a delayed translational increase in Arc levels (Seibt 2012). Cultured neurons have also been proposed to be similar to networks during sleep (Colombi 2016). Therefore, although our study is limited in that the chemical LTP protocol may not represent a physiological stimulus, the sustained late phase of Arc is still physiologically important to be studied as it is implicated in various mechanisms linked to memory consolidation.

It has been reported that Arc contains both a nuclear export signal (NES) as well as a nuclear localisation signal (NLS), suggesting that there are distinct mechanisms for Arc to shuttle in and out of the nucleus (Korb 2013). Although the mechanisms of regulation are not clearly established, previous work seems to suggest that dendritic and synaptic Arc occurs at an earlier time point than nuclear Arc. In this protocol, although we do note neuronal cytoplasmic Arc at the earlier time point (less than 4 hours after 4BF), we did not observe any distinct localisation to dendritic spines.

We have made use of time point assessment with immunohistochemistry to determine the spatio-temporal distribution of Arc after 4BF induction. We showed that Arc is expressed first in the neuronal cytoplasm and nucleus, and subsequently in astrocytes at a much longer time scale, and that Arc may be released from the neurons and subsequently taken up by surrounding astrocytes. A limitation of immuno-fluorescent staining, however, is that we can only have a snapshot of the expression profile at a given point in time. Given that neuronal networks have variability, the level of expression of Arc also has variability, so finer changes may not be able to be detected by taking a large sample and averaging (in order to determine significant levels of changes amidst the large variability). Alternative methods include using GFP fusion proteins and real time imaging. GFP fusion proteins, though, have other caveats such as issues of overexpression (Steward 2017), onset and offset kinetics, and also effects on post-transcriptional modifications and protein degradation. In particular, formation of structures like Arc capsids may be impeded by the fusion protein.

It is known that Arc can be induced by various protocols *in vitro* and *in vivo* with different temporal profiles of expression. It is likely that different pools of Arc exist and are induced and regulated differently. As pointed out in a review by Nikolaienko et al. (Nikolaienko 2018), Arc could potentially act as an organiser of long-term synaptic plasticity to mediate changes in connection strength in either direction, configuring the neuron to the context of ongoing neuronal activity. For this aspect, more studies should be done to investigate the regulation of spatiotemporal distribution of Arc.

### Implications of astrocytic Arc in memory

The pharmacological stimulation used here was able to robustly induce Arc, especially Arc in the nucleus, which makes this model an attractive one in the study of Arc dynamics, given that nuclear Arc is thought to be relevant for long-term memory consolidation. Here we reported an elevated expression of nuclear Arc at 4 hours, which corresponds to the time frame of late-LTP. Specifically, we report that this nuclear expression of Arc necessarily precedes that of the Arc level increase observed in astrocytes, showing that astrocytic Arc accumulation occurs at a later time point and as a consequence of neuronal activation and expression of Arc. This is in contrast with previous reports of Arc expression in astrocytes *in vivo*, where LTP was induced in the dentate gyrus by electrical stimulation of the perforant path (Rodriguez 2005). These studies found an increase in Arc expression in both neurons and astrocytes corresponding to the same time frame. This could be due to differences in the type of stimulation and the responses they elicit *in vivo* and *in vitro*, or the fact that our quantification methods were not sensitive enough to pick up small to moderate increases in astrocytic Arc at the early time points. It is clear from our cultures that at the later time point of 24 hours, there are elevated Arc levels, and this is mainly due to Arc within astrocytes. Such a phasic pattern of Arc expression could point to Arc’s role as a signalling molecule that carries information of neuronal activity to glial cells. Specifically, it could relay to the astrocytes the type of neuronal activation (in this case increased synchronised bursting). Modulation of astrocytic activity alone has been shown to be sufficient to induce LTP in the hippocampus (Adamsky 2018), and it is also known that the astrocyte-neuron lactate transport is essential for long term memory formation (Suzuki 2011). Astrocytes and neurons communicate bidirectionally and a range of neuromodulators can impact astrocytic activity, either directly or via regulation of specific gliotransmitter release (Covelo 2018). Therefore, it is possible that Arc could be a signal that communicates information regarding neuronal activation from neurons to astrocytes, potentially allowing astrocytes to participate in the subsequent interactions towards memory consolidation.

We have shown that Arc within astrocytes may associate with GFAP, though the significance of this is unknown. GFAP has been extensively studied in glial reactivity (Eng 1994), and could potentially play a role in astrocytic involvement of learning and memory (Choi 2016, Pearson-Leary 2016). Further elucidation of the interaction between Arc and GFAP may provide insights into how they might function to influence neuronal plasticity.

The activity-dependent accumulation of Arc in astrocytes could have important disease implications. Arc has been directly implicated in Alzheimer’s disease (AD) as Arc-endosomes are known to traffic APP to the plasma membrane to be cleaved to produce beta amyloid (Aβ) in an activity-dependent manner (Wu 2011). Aberrant expression of Arc was also found in mouse models of AD (Rudinskiy 2012). Astrocytes, on the other hand, are known to engulf plaques consisting of Aβ aggregates, and the integrity of this function is known to be crucial in the progression of AD (Gomez-Arboledas 2018).

### Accumulation of Arc in astrocytes

A number of immediate-early genes (IEGs), in particular *c-fos* and *c-jun*, have been reported previously to be expressed in astrocytes, although they are mostly linked to inflammatory pathways and reactive gliosis (Kato 1995, Rubio 1997). In line with this, *in vivo* subpopulations of astrocytes have been found to express c-fos during the development of multiple sclerosis in a mouse model, linking the expression of IEGs in astrocytes with the activation of the immune system. The expression of Arc in neurons has also been implicated in inflammatory processes, with studies indicating an increase in the number of Arc expressing neurons with the rise in number of activated microglia, especially during a learning episode, using a neuroinflammation animal model of chronic lipopolysaccharide infusion (Rosi 2005). However, no reports of astrocytic Arc expression have been linked to any proposed functions in inflammatory responses. We also did not note any changes in astrocyte morphology following the 4BF treatment and thus the astrocytic Arc observed at 24 hours seems unlikely to be an indicator of reactive gliosis.

We have demonstrated that Arc is not transcribed within astrocytes and can be transferred via conditioned media from neuronal cultures actively producing Arc. This mechanism of intercellular transfer is supported by recent structural studies of Arc, highlighting its homology with viral capsid proteins (Ashley 2018, Pastuzyn 2018, Erlendsson 2020). These studies were done in neuronal cultures lacking astrocytes, as astrocytes and other dividing cells were selectively eliminated with pharmacological agents. Thus, the dynamics of intercellular transfer of Arc capsids between neuron and glia remains unknown. The central nervous system is a highly dense environment with glia filling most of the spaces between neurons, especially in the hippocampus, where astrocytes are known to occupy discrete territories (Bushong 2002). The neuron-astrocyte ratio in our cultures is in the range of 40-60%, which resembles the distribution in the brain (Herculano-Houzel 2014). The density of our cultures was considerably high, such that intercellular communication is also more likely to occur between neurons and astrocytes.

We also assessed the localisation of astrocytic Arc and found that it does not associate with ALIX, a protein regulating endosomal trafficking and part of the extracellular vesicle (EV) components previously shown to mediate the intercellular transfer of Arc (Pastuzyn 2018). This could indicate that the astrocytic Arc we observed may not be part of the endocytic pathway. It is known that astrocytes resist HIV fusion, but can engulf material containing virus (Russell 2017), therefore the mechanism of entry for astrocytic Arc may not be simply based on EV transfer and could involve a more active role from the astrocytes.

Experiments with dynasore and/or other inhibitors of endocytosis could be further explored and optimised to determine if astrocytic Arc accumulation does indeed depend upon endocytosis. We have made use of a pretreatment protocol that was described in Pastuzyn et al. (Pastuzyn 2018), which was sufficient to block neuronal Arc capsid transfer. However, the induction protocol and the length of time of Arc expression used in this study differs from ours, and thus a similar pretreatment may not be enough to inhibit astrocytic endocytosis in our cultures. In another set of their experiments done on HEK cells over-expressing Arc, dynasore was used continuously in order to block intercellular transfer, showing that different inhibition protocols might need to be used depending on the culture and preparation. Besides clathrin- and caveolin-dependent endocytosis, astrocytes are also known to have a constitutive endocytic pathway that is dynamin independent and regulated by intracellular Ca^2+^ (Jiang 2009). Therefore, the inhibitory effect of dynasore may not act on this type of endocytosis. Furthermore, it is known that certain viruses such as simian virus 40 (SV40) can make use of dynamin-independent endocytic pathways for cell entry (Damm 2005). Since Arc potentially could function like viral capsids, the mechanism of entry of Arc into cells could also be distinct, depending on the cell type. This is therefore an important aspect to be further investigated.

We have not been able to show the presence of astrocytic Arc *in vivo* using the NEE behavioural protocol. This could be due to the low levels of Arc induced by the NEE, in contrast to highly abundant Arc in the cultures induced by 4BF. The relative low levels of Arc may therefore not be sufficient to induce astrocytic Arc accumulation. Alternative Arc induction methods which can lead to high level of Arc expression, such as fear conditioning or seizure protocols could be attempted. Rodriguez et. al. (Rodriguez 2005) reported the presence of astrocytic Arc after electrophysiological induction in anaesthetised rats (50 pulses at 250 Hz), though at a different timeline as what we report here. The expression dynamics therefore could depend on the type and intensity of the stimulus *in vivo* as well.

To determine the structure of Arc within astrocytes, super-resolution microscopy such as Stochastic Optical Reconstruction Microscopy (STORM) (Rust 2006) can be used. STORM has been used effectively to study capsid structures of viruses (e.g. (Pereira 2012, Laine 2015). It is also unknown if Arc would remain in capsid structures inside the astrocytes or only assume this form during the transfer. This could potentially be answered by comparing the macromolecular structures of Arc within astrocytes and extracellularly. If the accumulation of astrocytic Arc does take place via capsids from neurons with high level of Arc expression, the contiguous spread of Arc would be based on proximity. Given that our cultures are highly dense, where astrocytes are in contact with multiple neurons and other astrocytes, we did not note the effect of proximity. However, to investigate if Arc transfer is limited by distance, we could make use of less densely cultured cells or preparations of co-culture where neurons and astrocytes can be grown separately.

In conclusion, we established the spatiotemporal expression profile of Arc in mixed neuronal cultures after chemical LTP. We describe a distinct order of localisation, not only in neuronal subcellular locations but also the translocation of Arc from neurons to astrocytes. Our results suggest that Arc may be released from the neurons expressing high levels of Arc during periods of intense network activity and is subsequently taken up by surrounding glial cells. The existence of Arc within astrocytes, not originating from endogenous transcription, highlights the importance of characterising and studying its downstream functional role, especially in the context of learning and memory.

